# A conserved immune dysregulation is associated with infection severity, risk factors prior to infection, and treatment response

**DOI:** 10.1101/2025.04.18.649544

**Authors:** Ananthakrishnan Ganesan, Andrew R. Moore, Hong Zheng, Jiaying Toh, Michael Freedman, Andrew T. Magis, James R. Heath, Purvesh Khatri

## Abstract

Older age, being male, obesity, smoking, and comorbidities (e.g., diabetes, asthma) are associated with the increased risk for severe infections. We hypothesized that there is a conserved common immune dysregulation across these risk factors. We integrated single cell and bulk transcriptomic data and proteomic data from 12,026 blood samples across 68 cohorts to test this hypothesis. We found that our previously described 42-gene Severe-or-Mild (SoM) signature, is associated with each of these risk factors prior to infection. Furthermore, we found this conserved immune signature is modifiable using immunomodulatory drugs and lifestyle changes. The SoM score predicted patients with sepsis that would be harmed from hydrocortisone treatment, and patients with asthma that will not respond to monoclonal antibody treatment. Finally, the SoM score was associated with all-cause mortality. The SoM signature has the potential to redefine the immunologic framing of baseline immune state and response to chronic, subacute, and acute illnesses.

## Introduction

Older age [1–3], being male [2, 4–7], obesity [8–10], smoking [11, 12] and co-morbidities such as asthma [13, 14] and diabetes mellitus [8, 15, 16] have been identified as risk factors for severe outcomes in infectious diseases. More than 70% of US adults are estimated to have at least one risk factor for severe infection and over 40% have two or more risk factors [15, 16]. Notably, the risk for severe outcomes from infection increases with the number of risk factors, suggesting they are additive [17]. Despite the overwhelming epidemiological evidence, the underlying mechanisms for these risk factors have been largely investigated independent of each other. Importantly, how these risk factors affect the underlying immune responses prior to infection, and whether they affect the same immunological pathways that increase the risk of severe outcomes, including mortality, remains unknown.

Previously, we have described a 42-gene signature score, called Severe-or-Mild (SoM) score, which predicts at presentation (i.e., prior to outcome is known) which patients with viral infection will progress to have severe outcome with high accuracy [18]. Furthermore, we found that the SoM signature is evolutionarily conserved across at least 13 viruses in humans [18] and macaques [19]. Finally, we found that 42 genes in the SoM score are derived from four modules, which are divided into two detrimental and two protective modules, where higher expression of detrimental modules and lower expression of protective modules at presentation is associated with increased and decreased risk of severe outcomes, respectively. Since the 42 genes and the associated pathways are not specific to response to viral infections, we hypothesized that subjects with bacterial infections would also have a higher SoM score at presentation compared to healthy subjects. Given the strong association of the SoM score at presentation with the severity of infection, we hypothesized that the SoM score will be higher prior to infection in subjects with one or more risk factors for severe outcome compared to those without any risk factor.

To test these hypotheses, we carried out the largest multi-omics, multi-cohort analyses to date by integrating gene expression data from (1) patients with bacterial infection at presentation (17 studies, 1,295 samples), (2) patients with burn/trauma (30 subjects) or sepsis (176 subjects), (3) single-cell RNAseq (scRNA-seq) from patients with infection (bacterial or viral) (584,260 cells, 223 samples, 4 studies), (4) patients with asthma (4 studies, 603 subjects), (5) healthy subjects with a risk factor for severe infection (38 studies, 1,821 samples), (6) the Framingham Heart Study (5,321 samples), and (7) healthy obese subjects randomly assigned to calorie restricted diet (70 subjects). Further, we integrated proteomics profiles of 2,487 subjects with these gene expression data. Across these 12,026 individuals from 68 independent studies, our analyses found that the SoM score was associated not only with the severity of bacterial infection, asthma, and risk factors at presentation, but also their treatments, and was associated with all-cause mortality.

## Results

### The SoM host response signature score is conserved in bacterial infections and is associated with severity

We first investigated whether the SoM score, which was identified using only respiratory viral infections [18, 20], was also correlated with severity in patients with bacterial infection. We identified, curated, and processed 17 datasets [21–37] that profiled 1,295 blood samples from 467 healthy controls and 828 patients with bacterial infections with varying level of severity (Figure 1A and Table S1; Methods). Next, we calculated the SoM score for each sample as described before [18]. Briefly, for each module in each sample, we computed a module score as the geometric mean of the expression of genes in the module. Next, we computed the detrimental score as the sum of scores of modules 1 and 2, and protective score as the sum of scores of modules 3 and 4. Finally, we computed the SoM score for a sample as the difference between the detrimental and protective scores, which represents the dysregulation within the immune system such that higher SoM score corresponds to higher immune dysregulation.

**Fig. 1.**
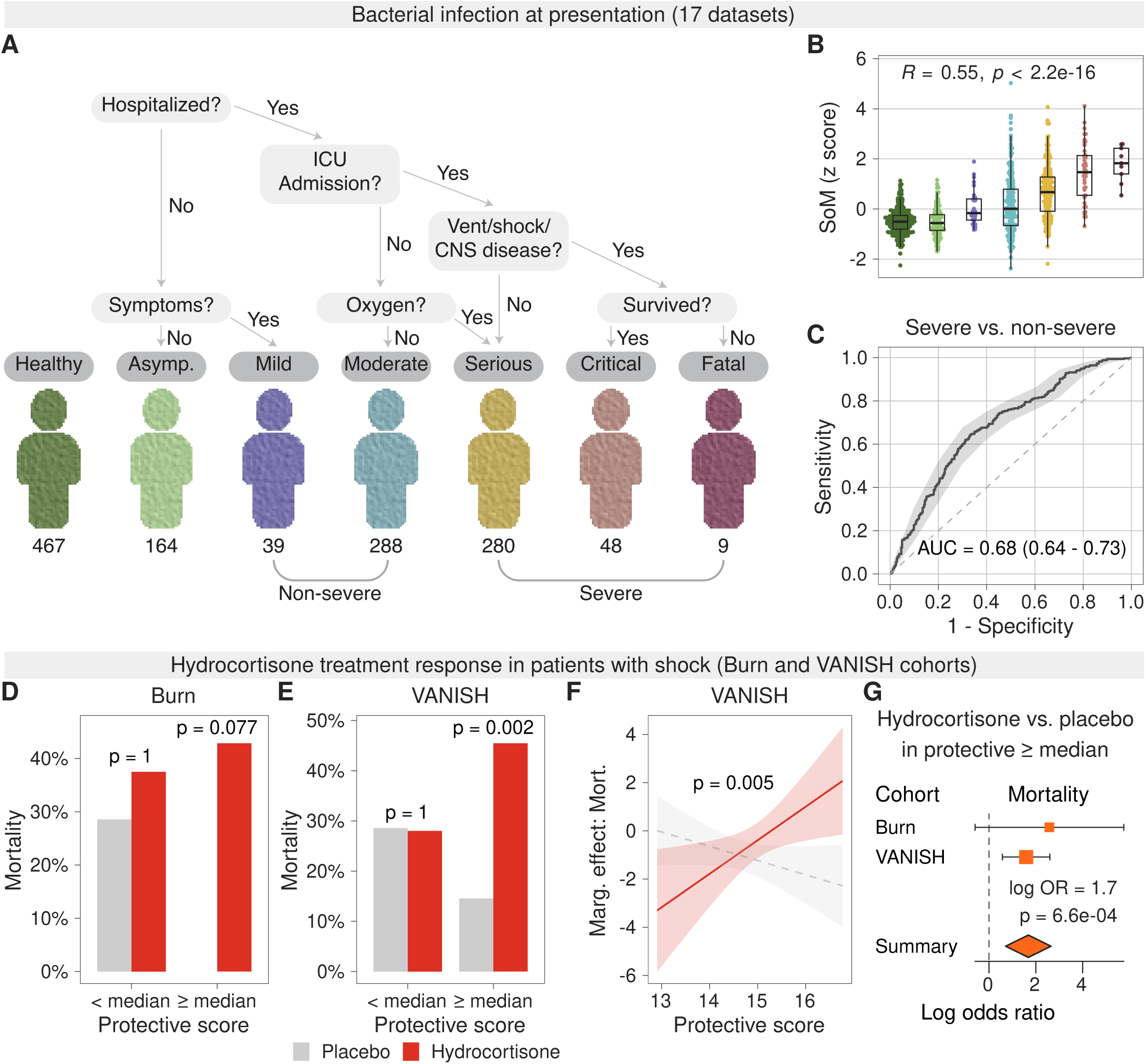
(**A**) Criteria for assigning bacterial infection severity categories to samples and the number of samples in each severity category. Mild and moderate categories were considered non-severe and serious, critical, and fatal categories were considered severe. (**B**) Variation of the z-scaled SoM score with severity of bacterial infection. R denotes Pearson correlation, and p its corresponding p-value. (**C**) Receiver operating characteristic curve for discriminating severe infection from non-severe bacterial infection using the SoM score. AUC denotes area under curve. 95% confidence interval estimates for AUC are shown alongside the AUC. (**D-E**) Bar plot of mortality comparing placebo and hydrocortisone arms for subjects with baseline protective scores lower than and higher than median in the (**D**) Burn and (**E**) Vanish cohorts. P-values were calculated using the Fisher test. (**F**) Interaction plot showing the marginal dependence of mortality on the protective score for the placebo and hydrocortisone arms in the Vanish cohort. Logistic model p-value for the interaction term between protective score and steroid arm is shown. **G**) Forest plot depicting log odds ratio for mortality among patients with protective score greater than median in the Burn and VANISH cohorts. Summary effect size and p-value were computed using a mixed effects model.

The SoM score was positively correlated with severity of bacterial infections (r=0.55, p<2.2e-16; Figure 1B). Similar to viral infections, the detrimental module scores were positively (r=0.61, p<2.2e-16; Figure S1A), and the protective module scores were negatively correlated with severity (r=-0.16, p=1.1e-08; Figure S1B). Finally, the SoM score distinguished opatients with severe bacterial infection from those with non-severe bacterial infection with an area under the receiver operating characteristic (AUROC) curve of 0.68 (95% CI: 0.64-0.73; Figure 1C). Collectively, these results strongly suggest that in patients with severe infection, irrespective of bacterial or viral etiology, there is a significant overlap in dysregulation of host response.

### Protective host response scores are associated with differential response to hydrocortisone

Over the last 5 decades, more than 100 large clinical trials in sepsis have failed due to patient heterogeneity [38]. Among the multiple reasons for these failures, one of the reasons is the variability in immune responses, such that potential treatment benefits in patient subgroups are negated by harms in others. We hypothesized that quantifying detrimental and protective responses in sepsis patients will reduce the heterogeneity to allow identification of patients who will benefit or be harmed by immunomodulatory treatments.

To test this hypothesis, we first investigated whether the SoM score was associated with increased mortality in two independent cohorts, which included (1) burn patients (n=30) with vasodilatory shock (Burn cohort; GSE77791) [39, 40] that were randomized to receive hydrocortisone versus placebo and (2) the VANISH [41] trial, which included 176 patients with septic shock, of which 117 were randomized to either hydrocortisone or placebo. Across both cohorts, high SoM score was significantly (p=0.035) associated with increased mortality at baseline (Figure S1C). Next, we investigated whether steroid treatment affects protective or detrimental scores. In the Burn cohort, steroid treatment significantly reduced the protective scores (p=0.004; Figure S1D) on day 7 compared to placebo, but not the detrimental scores, which increased in both groups (p>0.05; Figure S1E). In other words, steroid treatment reduced the protective immune responses significantly more than the detrimental response, which in turn increased the immune dysregulation within a patient. Therefore, we hypothesized that steroid treatment would harm patients with higher protective scores at baseline as it would increase the immune dysregulation. To test this hypothesis, we divided the Burn cohort patients in “high protective” and “low protective” score groups using the median of the protective scores as a threshold. In the high protective score group, the 28-day mortality was 43% in patients who were treated with hydrocortisone compared to 0% in patients who received placebo (Figure 1D), although this difference was not statistically significant (p=0.077), possibly due to small sample size.

Next, we used the VANISH [41] trial to replicate this result. We evaluated the effect of hydrocortisone on 28-day mortality in these patients with septic shock. Similar to the Burn cohort, we divided these patients into “high protective” and “low protective” score groups using the median of the protective scores. Similar to the Burn cohort, there was no difference in mortality in the low protective score group (p=1; Figure 1E), validating the results in the Burn cohort. However, in the high protective score group, the 28-day mortality was significantly higher for those treated with steroid than the placebo (45.5% vs. 14.5%, odds ratio (OR)=4.79, 95% CI: 1.59-15.55, p=0.002; Figure 1E). The results remained the same when considering only 117 patients that were randomized (50% vs. 20%, odds ratio=3.74, 95% CI: 1.1-14.6, p=0.03; Figure S1F). Logistic regression analysis, controlling for sex and the interaction with APACHE II score, to evaluate the interaction between steroid treatment and the protective score found a significant interaction indicating that patients with high protective scores (i.e., low-risk patients), and treated with hydrocortisone, experienced a higher rate of 28-day mortality (p=0.005; Figure 1F). Similar analysis using only the detrimental scores found no such interaction and difference in mortality by treatment (Figure S1G-H). Across both datasets, hydrocortisone treatment in high protective score group was associated with significantly increased risk of mortality (summary log OR=1.68, 95% CI: 0.71-2.65, p=6.6e-04; Figure 1G). Together, these results demonstrated that the SoM score, along with its constituent detrimental and protective scores, is also conserved in patients with bacterial infections, associated with severity of their outcomes, is differentially modulated by steroid treatment, and could be used to identify patients who may be harmed by steroid treatment.

### Detrimental host response modules are primarily driven by neutrophils

Next, we explored the immune cell types associated with each of the four gene modules in the SoM signature. We collected, curated, and integrated four publicly available whole blood single-cell transcriptomic datasets (584,260 cells, 223 samples) [42–45] from healthy subjects and patients with bacterial or viral infection of varying degrees of severity (Figure 2A-C, Methods). We chose these datasets as they included scRNA-seq data from neutrophils. Among the detrimental modules, module 1 score was the highest in mature neutrophils, whereas module 2 score was almost exclusively expressed in immature neutrophils (Figure 2D). Among the protective modules, module 3 score was the highest in monocytes but also detected in T and NK cells, whereas module 4 score was primarily expressed in T and NK cells (Figures 2D-E, Figure S2). These results were in line with our previous observations, where we found that several genes in the detrimental modules were known to be expressed in mature and developing neutrophils, whereas the genes in the protective modules were known to be expressed in monocytes, T, and NK cells [46]. Together, these results suggest the detrimental host responses during acute infections are primarily driven by different neutrophil subsets, whereas protective responses are driven by monocytes, T, and NK cells. These results demonstrate that the SoM score quantifies immune dysregulation at a system-level by integrating gene modules from both myeloid and lymphoid immune cells.

**Fig. 2.**
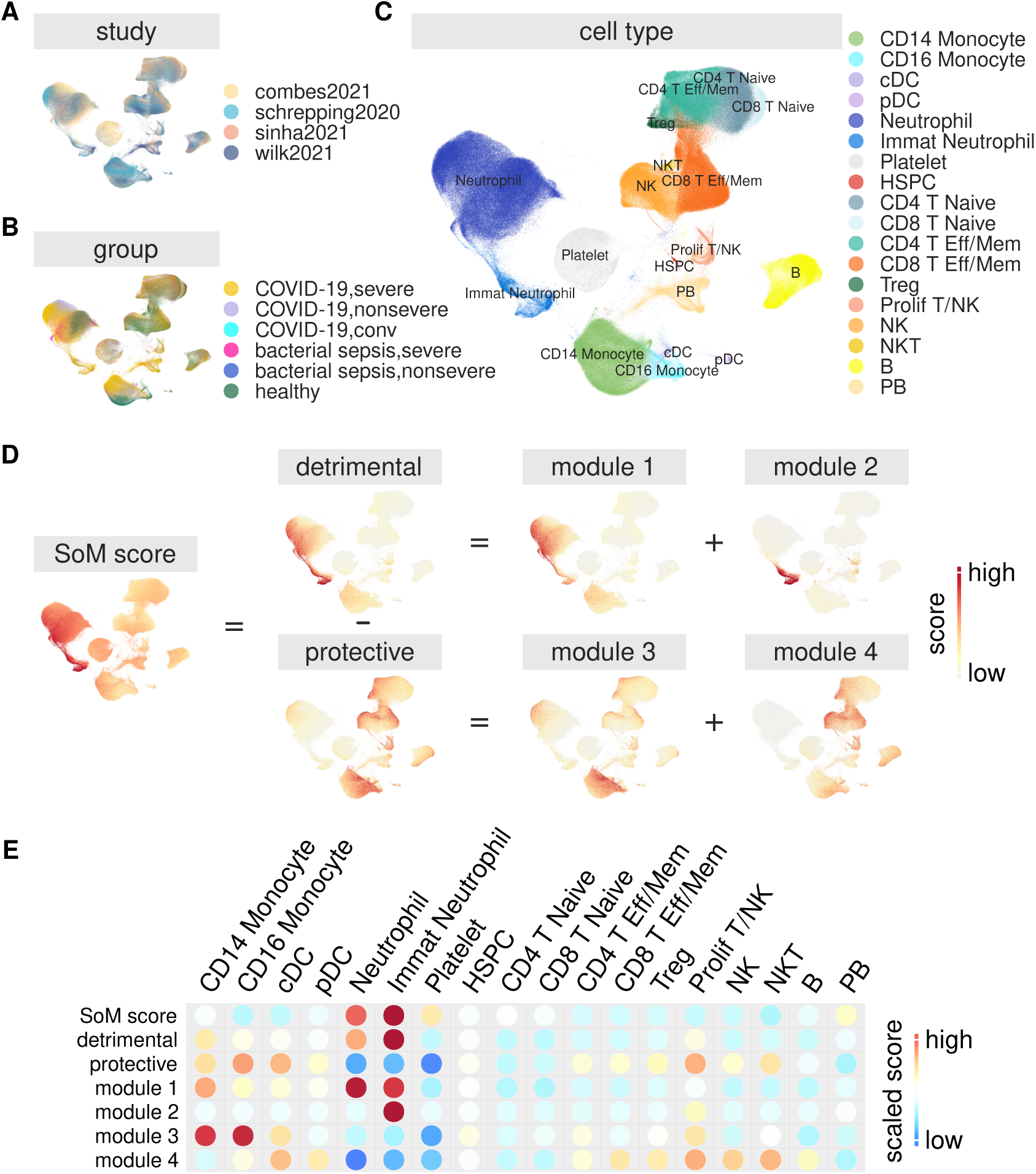
(**A-D**) UMAP visualization of single cell whole blood transcriptome data colored by (**A**) dataset, (**B**) phenotype, (**C**) cell type, and (**D**) SoM, detrimental, protective, and modules 1-4 scores. Each point is a cell and UMAP was projected based on the gene expression for each cell. (**E**) Heatmap of average score value for the 7 scores in 20 cell types.

### Immune dysregulation is higher in patients with severe asthma and associated with poor response to monoclonal antibody treatment

Respiratory viral and bacterial infections have been repeatedly associated with and linked to airway remodeling and severe asthma [47, 48]. Given that the SoM score was associated with severity in both viral and bacterial infections, we investigated whether patients with asthma, who tend to have severe exacerbations, also had a higher SoM score compared to healthy subjects before the exacerbations. To this end, we used 498 transcriptome profiles (87 healthy, 77 moderate asthma, 334 severe asthma) from U-BIOPRED (GSE69683) [49], a multicenter prospective cohort study across 16 clinical centers in 11 European countries. The SoM score was significantly higher in patients with the severe asthma compared to healthy controls and those with moderate asthma (JT trend test p-value<2.2e-16; Figure 3A), which was driven by significant increase in the detrimental response (JT trend test p-value<2.2e-16; Figure 3B) and decrease in the protective response (JT trend test p-value=3.7e-07; Figure 3C). While the increase in detrimental response was driven by increase in both detrimental modules 1 and 2 (JT trend p-values of 1.7e-14 and 7.7e-12, respectively; Figure S3A), the decrease in protective response was solely due to reduced module 4 scores with increased severity (JT trend test p-value=2.1e-09; Figure S3A) with no significant change in module 3 (JT trend test p-value>0.1; Figure S3A).

**Fig. 3.**
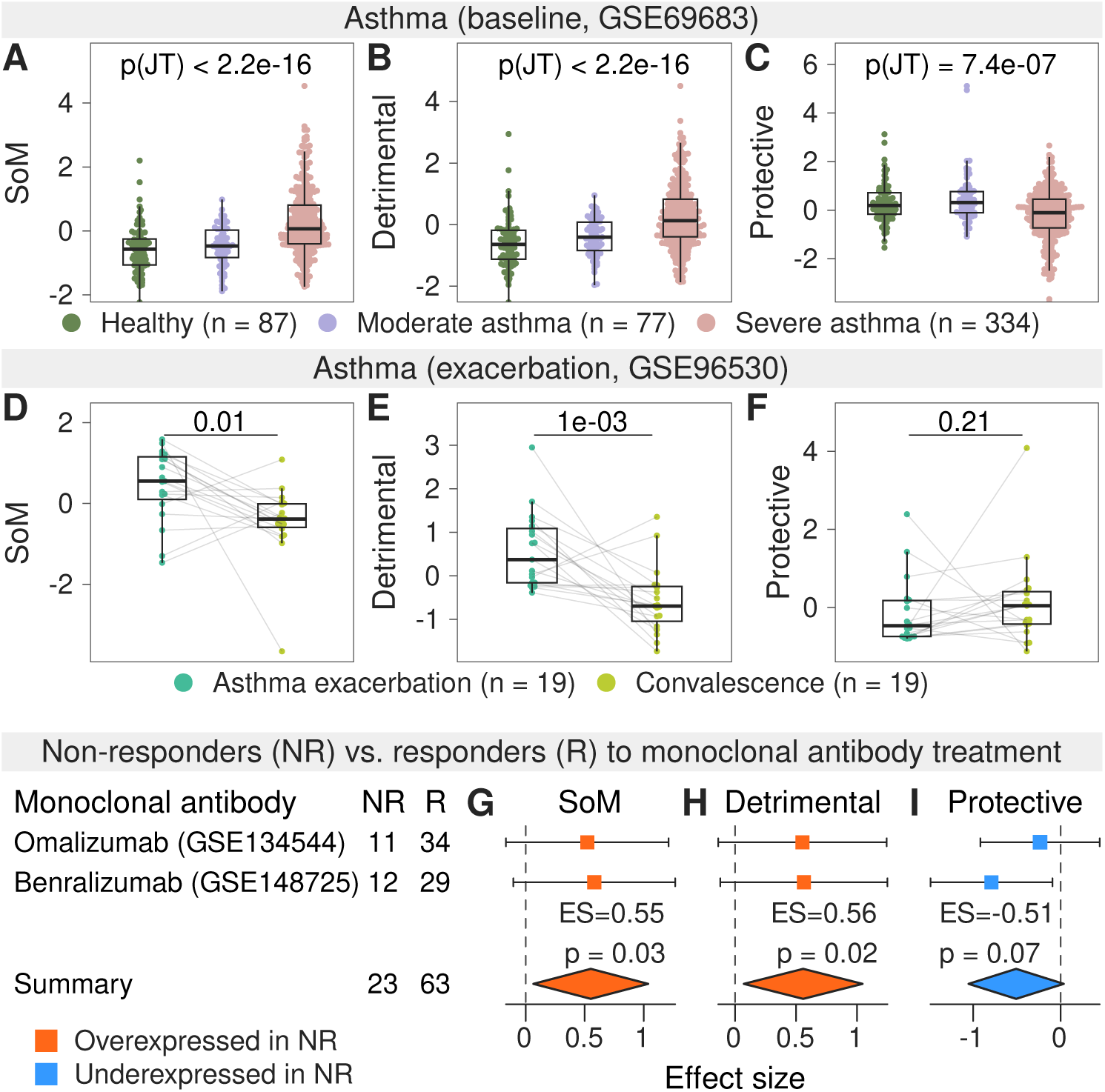
(**A-C**) Variation of the z-scaled (**A**) SoM, (**B**) detrimental, and (**C**) protective scores with asthma severity. P-values were calculated sing the Jonckheere–Terpstra trend test. (**D-F**) Comparison of the z-scaled (**D**) SoM, (**E**) detrimental, and (**F**) protective scores at asthma exacerbation and after convalescence. P-values were calculated using the paired Wilcox test. (**G-I**) Forest plots of dataset-wise and summary effect sizes for the (**G**) SoM, (**H**) detrimental, and (**I**) protective scores in non-responders compared to tresponders for monoclonal antibody reatment for asthma. P-values for the summary effect sizes are shown. NR denotes non-responders and R responders.

Next, we asked whether there was further increase in the SoM scores during asthma exacerbation. In a cohort of 19 patients with asthma, profiled during and after asthma exacerbation (GSE96530) [50], the SoM score was significantly higher in peripheral blood during exacerbation than convalescence (paired Wilcox test p-value=0.011; Figure 3D), which was driven by a significantly higher detrimental score (paired Wilcox test p-value=0.0012; Figure 3E) without significant change in the protective score (paired Wilcox test p-value>0.2; Figure 3F) during exacerbation. Finally, similar to U-BIOPRED, higher SoM score during exacerbation was driven by significantly higher detrimental modules 1 and 2 during exacerbation (paired Wilcox test p-value*≤*0.011; Figure S3B) and significantly lower protective module 4 (paired Wilcox test p-value=0.02; Figure S3B).

Patients with neutrophilic asthma are known to respond poorly to monoclonal antibody (mAb)-based treatments [51, 52]. Because the detrimental modules were primarily expressed in neutrophils, which were higher in those who tend to have severe asthma, we investigated whether the detrimental scores are significantly higher in patients with asthma that do not respond to mAb treatment. To test this hypothesis, we used transcriptome data from two cohorts that profiled asthma patients treated with either omalizumab (GSE134544) [53] or benralizumab (GSE148725) [54]. GSE134544 performed transcriptome profiling of whole blood from 45 patients with moderate-to-severe asthma treated with omalizumab (34 responders, 11 non-responders) [53]. GSE148725 performed transcriptome profiling of whole blood from 41 patients with severe eosinophilic asthma treated with benralizumab (29 responders, 12 non-responders) [54]. Our analysis only included samples prior to treatment initiation. Across both cohorts, non-responders had significantly higher SoM score (ES=0.55, p=0.03; Figure 3G) that was driven by significantly higher detrimental score (ES=0.56, p=0.02; Figure 3H), prior to start of mAb treatment. Although non-responders had lower protective score, it was only marginally statistically significant (ES= −0.51, p=0.07; Figure 3I).

Collectively, our results suggest that the immune dysregulation, as quantified by the SoM score, is higher in patients with severe asthma than those with mild/moderate asthma prior to exacerbation, increases further during asthma exacerbation, and is higher in patients with asthma that do not response to mAb treatment prior to treatment initiation.

### Detrimental host response modules are significantly associated with risk factors for severe infections prior to infection

Because higher SoM score was associated with severe asthma and with severe outcomes in infectious diseases, irrespective of bacterial or viral infections, we next asked whether it was also significantly associated with one or more risk factors for severe infections *prior* to infection. To answer this question, we identified, curated, and preprocessed 38 case-control studies comprising 1,821 blood samples (Figure 4A and Table S2; Methods). These studies included (1) eight datasets comparing healthy 60 years or older subjects (n=249) with those 40 years or younger (n=126) [55–59], (2) sixteen datasets comparing healthy males (n=443) with healthy females (n=510) [60–73], (3) four datasets comparing otherwise healthy subjects with BMI*≥*30 (n=95) with those with BMI*≤*25 (n=85) [74–77], and (4) 10 datasets comparing patients with type 1 or type 2 diabetes (n=185) with healthy controls (n=128) [78–85]. None of these subjects had any known infections.

**Fig. 4.**
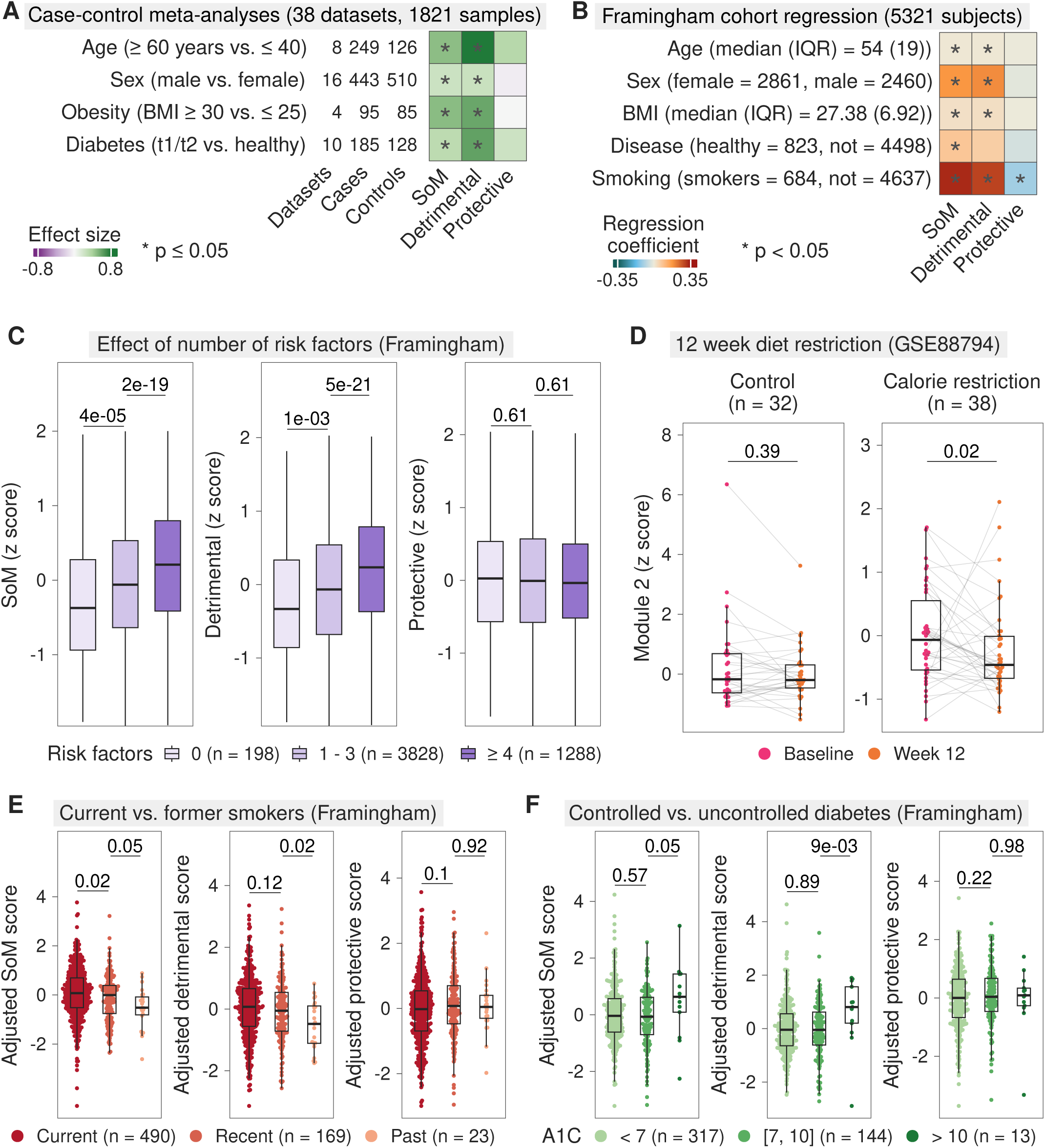
(**A**) Summary of case-control meta-analyses for risk factors of infection. Heatmap depicts summary effect size for a given score and risk factor. * indicates p*≤*0.05, where p is the p-value for the summary effect size from the meta analysis. (**B**) Summary of linear regression models for risk factors of infection in the Framingham cohort. Heatmap depicts the regression coefficient for a given score and risk factor. * denotes p<0.05, where p is the p-value for the regression coefficients. (**C**) Variation of the z-scaled SoM, detrimental, and protective scores with number of risk factors. P-values were computed using the Wilcox test. (**D**) Comparison of the z-scaled module 2 score at baseline and after 12 weeks for the control and calorie restriction groups in a randomized diet restriction study. P-values were computed using the paired Wilcox test. (**E-F**) Variation in the SoM, detrimental, and protective scores adjusted for age, sex, BMI and (**E**) disease or (**F**) smoking status among (**E**) current, recent (quit within the last 5 years), and past (quit more than 5 years ago) smokers and (**F**) subjects with controlled (A1C<7), moderately uncontrolled (7*≤*A1C*≤*10), and highly uncontrolled (A1C>10) diabetes. P-values were computed using the Wilcox test.

SoM score was significantly higher (p*≤*0.05) in older subjects compared to those younger than 40, in males compared to females, in those with BMI*≥*30 compared to those with BMI*≤*25, and in patients with diabetes compared to healthy people (Figure 4A). The higher SoM score was primarily driven by an increase in the detrimental score that was signifiantly higher in subjects with the risk factors, whereas the protective score was not significantly different (Figure 4A).

We could not analyze the impact of presence of more than one risk factors (e.g., an older man with BMI*≥*30) in these case-control studies. However, otherwise healthy people can have more than one risk factors (e.g., older male with BMI>30 who smokes). Furthermore, the studies included in our analysis only compared tails of a given risk factor (e.g., *≤*40 years old vs. *≥*60 years old, BMI*≤*25 vs. BMI*≥*30, etc.). In other words, these studies were not representative of the heterogeneity in the real-world population. To address these limitations, we used whole blood transcriptome profiles of 5,321 subjects in the Framingham Heart Study (Figure 4B) [86]. Using clinical questionnaires and summaries of medical records, we collected age, sex, BMI, smoking status, and disease status for each subject (Figure 4B; Methods). The Framingham Heart Study had two advantages over the case-control studies: (1) it allowed analysis over the entire range of values for a risk factor, and (2) allowed analysis when individuals had more than one risk factors.

We used linear regression to investigate whether any of the risk factors were significantly associated with the SoM, detrimental, and protective scores. We used age and BMI as continuous, and sex, smoking status, and disease status as binary independent variables. All risk factors (age, sex, BMI, presence of disease, smoking) were significantly positively associated with the SoM score in the Framingham cohort (p<0.05, Figure 4B). Similar to the case-control studies, the positive association of the SoM score with the risk factors was driven by significant positive association of the detrimental score with age, sex, BMI, and smoking status (p<0.05, Figure 4B). The protective score was not associated with any risk factors, except it was significantly lower in those who smoked (Figure 4B). Notably, although neither detrimental nor protective score were significantly associated with a presence of disease, the SoM score was significantly associated with the presence of a disease. Overall, these results were consistent with the results from the case-control meta-analyses. Together, these results suggests that the immune dysregulation, represented by the SoM score, is common across several known risk factors for severe outcomes in infectious diseases prior to infection. Furthermore, these results showed that the increased immune dysregulation is primarily driven by increase in the detrimental scores without the corresponding increase in the protective scores. More importantly, our results demonstrate the presence of a conserved common immune dysregulation across multiple risk factors for severe outcome in infectious diseases.

### Detrimental host response increases with the number of risk factors

Epidemiological data have shown that the risk for severe outcomes from infection is increases with the number of risk factors in an individual [17]. Therefore, we investigated whether the SoM, detrimental, and protective scores are correlated with the number of risk factors present in an individual. We compared each of the three scores against the total number of risk factors present in each subject—BMI*≥*30, *≥*60 years old, male, current or former smoker, and each disease were counted as one risk factor. Subjects ranged from having no risk factors to as many as 8 risk factors. The SoM and detrimental scores significantly correlated with the number of risk factors in an individual such that they were highest in subjects with 4 or more risk factors, and lowest in subjects with no risk factors (Figure 4C, Wilcox p<0.05). In contrast, the protective score was not correlated with the number of risk factors in an individual (Figure 4C, Wilcox p>0.1). These results further supported our analysis that the SoM score represents a conserved common immune dysregulation independently associated with risk factors, and is additive as the number of risk factors in an individual increases.

### Immune dysregulation decreases with the management of risk factors

Our analysis of patients treated with hydrocortisone demonstrated that the detrimental and protective scores are modifiable. Because our analysis found that BMI at the time of gene expression was associated with the extent of immune dysregulation, we hypothesized that immune dysregulation (i.e., the SoM score) would decrease with lifestyle changes uch as reduced BMI, quit smoking, or controlling A1C in patients with diabetes.

To test this hypothesis, we used peripheral blood mononuclear cell (PBMC) transcriptome data from 70 healthy, overweight subjects (GSE88794) [87], of which 38 subjects were randomly assigned to a 20% calorie restriction diet for 12 weeks (calorie restriction group) and 32 were not (control group). The calorie restriction group had a mean weight loss of 5.6kgs over 12 weeks [87]. In this randomized study, module 2 decreased significantly in the calorie restriction group (paired Wilcox test p-value=0.02; Figure 4D), whereas it remained unchanged in the control group (paired Wilcox test p-value>0.05; Figure 4D). There was no significant difference in the SoM and module 1 scores in both the control and calorie restriction groups (paired Wilcox test p-value>0.05; Figure S4A-B), possibly due to neutrophils, which we found to drive module 1 (Figure 2), being filtered out in the PBMC samples.

Next, we investigated if the SoM score was lower among subjects that quit smoking. In the Framingham Heart Study, we identified 490 current smokers, 169 recent smokers who quit smoking within the last 5 years, and 23 who quit smoking at least 5 years prior (past smokers) to gene expression measurement in the Framingham cohort. The SoM score, adjusted for age, sex, BMI, and disease status, was the highest in current smokers and lowest in past smokers (p*≤*0.05; Figure 4E). The detrimental and protective scores were not different between current and recent smokers (p>0.1; Figure 4E), suggesting it took more than 5 years since quitting smoking for desired changes in immune dysregulation.

Finally, we investigated whether the SoM score was lower in patients with diabetes with lower A1C values. Among 474 patients with diabetes in the Framingham cohort, we identified 317 with controlled diabetes (A1C<7), 144 with moderately uncontrolled diabetes (7*≤*A1C*≤*10), and 13 with uncontrolled diabetes (A1C>10). After adjusting for age, sex, BMI, and smoking status, there was no difference in the SoM score between patients with controlled or moderately uncontrolled diabetes (p>0.05; Figure 4F), but was significantly higher in patients with uncontrolled diabetes compared to moderately uncontrolled diabetes (p=0.05; Figure 4F). This increase in the SoM score in patients with uncontrolled diabetes was driven by a significant increase in their detrimental score (p=0.009), without significant change in the protective score (Figure 4F). These results suggests patients with diabetes with lower A1C values have reduced immune dysregulation as represented by the SoM score.

Next, we compared the SoM score to the immune health metric (IHM) score [88], a 150-gene signature that delineated healthy individuals from those with monogenic diseases and correlated with aging in healthy individuals. None of the 42 genes defining the SoM score overlapped with the 150-gene IHM signature. Despite this, the SoM score and IHM score were inversely correlated in the Framingham cohort (R=-0.43, p<0.01; Figure S4C), suggesting a partial overlap of pathways between these two scores. This is not surprising since both were associated with age. However, there was a significant difference between the two signatures. Unlike the SoM score that is modifiable with intervention, the IHM score did not change significantly following 12-week calorie restriction (p>0.05; Figure S4D). Similarly, there was no difference in the IHM score between current, recent, and past smokers (Figure S4E), and in those with controlled or uncontrolled diabetes (Figure S4F).

### Increased abundance of proteins in modules 1 and 2 among subjects with risk factors

Next, we investigated whether these transcriptomics changes were also observed at the protein level using data from a cohort of 2,487 subjects from the Institute for Systems Biology (Arivale cohort) [89]. This data contained protein measurements in plasma corresponding to 5 of the 42 genes defining the SoM score (ADM and GRN from module 1; CEACAM8, AZU1, and OLR1 from module 2). Proteins from module 2 were positively associated with age, sex, BMI, and disease (Figure S4G). GRN from module 1 was positively associated with age, BMI, and disease. Each of the 5 proteins were significantly and positively associated with BMI (p<0.05; Methods; Figure S4G). The effect of sex was mixed, with men having higher abundance of proteins in module 2 compared to women but lower abundance of proteins in module 1 (p<0.05; Figure S4G). Overall, the results from analysis of proteomic data were mostly consistent with our observations from transcriptomic data.

### Higher risk of mortality in subjects with increased dysregulation of host response

Finally, we investigated whether immune dysregulation, as quantified by the SoM score, is associated with higher risk of all-cause mortality. In the Framingham cohort, 262 of the 5321 subjects died within 7 years after gene expression was profiled. We found age, sex, disease status, and smoking status were significantly associated with all-cause mortality in the Framingham cohort (Figure 5A). Importantly, the SoM score was independently associated with increased risk of mortality (HR=1.64, p=4.6e-05; Figure 5A). Therefore, we adjusted the SoM score by regressing out the effects of age, sex, BMI, disease and smoking status. A Cox regression model using the adjusted SoM score showed that it was significantly ssociated with the increased risk of mortality (HR=1.8, p=7.2e-06; Figure 5B). Next, when we divided the Framingham cohort into two groups using the median adjusted SoM score as “high SoM” and “low SoM” groups, those in the high SoM group had a significantly higher risk of all-cause mortality than those in the low SoM group (HR=1.36, p=0.013; Figure 5C). The adjusted SoM remained an independent risk factor of increased mortality even when considering mortality over a 10-year period (HR=1.44, p=3.3e-04; Figure S5A) or only those over 50 years of age (HR=1.73, p=2.3e-05; Figure S5B). This result strongly suggests that those with higher immune dysregulation than expected for a given age, sex, BMI, disease status, and smoking status are at a higher risk of mortality.

**Fig. 5.**
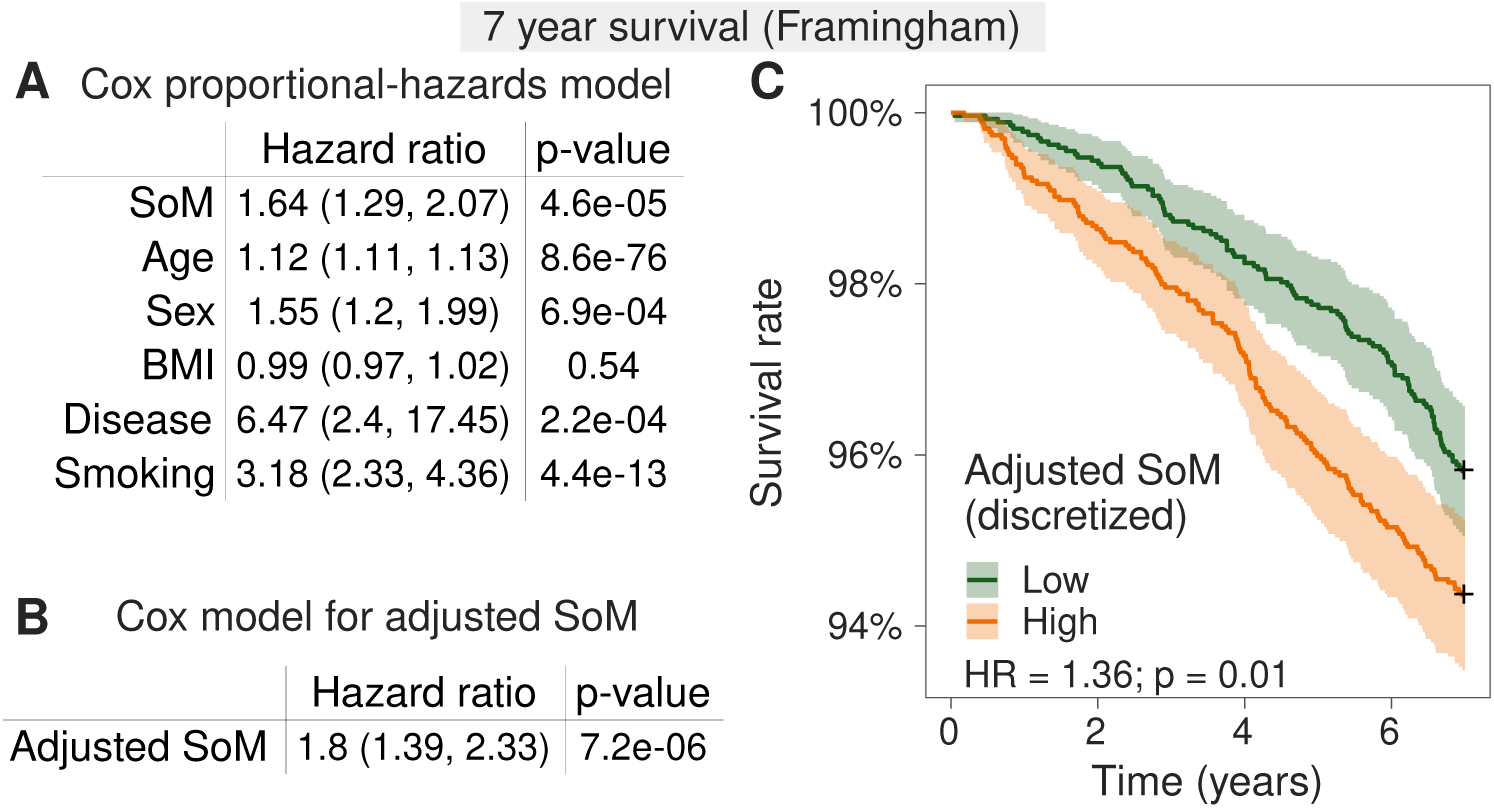
(**A-B**) Summary of Cox proportional-hazards models in the Framingham cohort evaluating the association with 7-year mortality for **A**) the SoM score, age, sex, BMI, disease and smoking statuses, and (**B**) the SoM score adjusted for age, sex, BMI, disease and smoking statuses. Hazard ratio, its 95% confidence estimates, and the corresponding p-values are shown. (**C**) Kaplan-Meier survival curve for the adjusted SoM score divided into high and low groups based on median value. Cox model hazard ratio and p-value are shown.

## Discussion

Several risk factors, including age, sex, BMI, smoking, and comorbidities such as diabetes and asthma, are associated with poor outcomes in patients with infectious diseases. Although these risk factors are known to alter immune status of an individual, it is unclear whether these risk factors lead to the same conversed immune responses. We hypothesized that these distinct risk factors associated with severe outcomes lead to a conserved immune status that would be associated with the increased risk of severe outcomes. Using more than 12,000 blood transcriptome profiles across 68 independent studies, we comprehensively tested this hypothesis using our previously described 42-gene signature in blood, referred to as the SoM signature [18] that predicts the risk of severe outcome at presentation. Although the SoM signature was identified and validated in only viral infections, we found it is also associated with severe outcomes in patients with bacterial infections. Strikingly, we found the SoM signature score was higher in those with any of the risk factors for severe outcomes, was positively correlated with the number of risk factors in an individual, and predicted all-cause mortality, even when they were not infected and despite the real-world biological, clinical, and technical heterogeneity cross 68 independent studies. Most importantly, we found that this common dysregulated immune response, represented and quantified by the SoM score, could predict response to steroids in patients with a critical illness such sepsis or burn, or to monoclonal antibody treatment in patients with asthma. Finally, we found diet restrictions and smoking cessation would also reduce this dysregulated immune response. Collectively, these results strongly support our hypothesis and demonstrate a presence of a dysregulated immune response that is conserved prior to and during infectious diseases, associated with severe outcomes, including all-cause mortality, and is modifiable through drug treatment and lifestyle changes.

We found moderate correlation between the 42-gene SoM score and the 150-gene IHM score [88], although none of the genes overlapped, suggesting some of the constituents of both signatures may be similar. For instance, one of the protective response modules, module 4, is preferentially expressed in NK cells, which were also a key constituent of the IHM. Despite this correlation and common immune constituents, our results also suggest that these two signatures represent distinct immune responses. Specifically, the IHM signature was identified using 22 monogenic immune diseases, whereas the SoM signature was identified from response to respiratory viral infections. This difference in how both signatures were identified suggest that the IHM signature is rooted in genetics, whereas the SoM signature is triggered by epigenetics and lifestyle choices. This is further supported by the fact that the SoM signature was modifiable through drug treatment and changes in lifestyle (reduced calorie intakes, smoking cessation), but the IHM signature was not as drug treatments or lifestyle changes will not change genetics of an individual. Future studies that integrate both the IHM and the SoM signatures have the potential to disentangle and quantify immune health status of an individual with which they are born and how it changes over their lifetime due to environmental exposures and lifestyle choices.

Our scRNA-seq analysis found that the four gene modules in the SoM signature are preferentially expressed in different immune cell types, suggesting that the SoM score quantifies immune dysregulation at a system level. Specifically, each of the four modules in the SoM signature are preferentially expressed in different immune cell types, which include both myeloid and lymphoid cells such that mature and immature neutrophils expressed the detrimental response modules and monocytes, T and NK cells expressed the protective response modules. It is likely that these four modules are further interacting with each other. For example, the detrimental modules include *BCL2L11*, *CEACAM8*, and *ORM1*, which are associated with lymphocyte apoptosis and immunosuppression [90–92]. Indeed, we found that the overall protective immune response in non-infected subjects with one or more risk factors was not significantly different compared to those without any risk factors. However, in patients with asthma or bacterial infection, one of the protective modules, module 4 that is preferentially expressed in T and NK cells, was significantly lower compared to healthy subjects. These suggest the immunosuppressive functions of these neutrophils, particularly the downregulation of lymphocytes, as a potential mechanism explaining the observed detrimental effect. Further studies are required to understand if and how the neutrophil-driven detrimental host responses are associated with lymphocyte-drive protective responses.

While the association between neutrophils and asthma severity is known [51], our results expand our understanding. Our analysis suggests this association is driven by specific subsets of neutrophils that express detrimental immune response modules. Our analysis also suggests that these neutrophil subsets are unlikely to be only associated with neutrophilic asthma as we found patients with moderate-to-severe asthma who do not respond to omalizumab and those with severe eosinophilic asthma who do not respond to benralizumab had higher detrimental scores, which are driven primarily by a subset of neutrophils. Future studies should focus on elucidating the role of the module 2 expressing immature neutrophils in asthma [93].

Heterogeneity of treatment effect in patients with sepsis, burn, or asthma has been a significant roadblock in advancing novel treatments for these diseases. Our results in using the SoM score to identify patients who may be harmed from hydrocortisone treatment or patients with asthma that do not respond to monoclonal antibody treatment suggests its potential utility for clinical trials for enrollment through predictive enrichment. Across three inflammatory diseases, using baseline SoM score (i.e., prior to treatment initiation), we showed that patients have differential response to immune-modulating therapies. These findings are in line with prior studies that have shown the association of immune endotypes with treatment responsiveness [41, 94, 95], but now provide insights into underlying conserved immune dysregulation across biological, clinical, and technical heterogeneity. In other words, by identifying a set of patients that may be harmed by or benefit from a potential treatment prior to treatment initiation, the SoM score has the potential to reduce the heterogeneity of treatment effect in future studies.

The SoM score had lower accuracy for predicting severity in patients with bacterial infection than those with viral infection. The reduced accuracy is likely due to interferon stimulated genes in module 3 that are more responsive to viral infection than bacterial infection. Nevertheless, its ability to identify which patients will be harmed by steroids in burn and bacterial sepsis patients suggest that while it may not be relevant to response to bacterial pathogens, it is important in identifying which patients should or should not be treated with steroids.

Most importantly, the SoM represents a common dysregulated immune response that is higher in individuals with risk factors prior to infection, predicts severity of infection at presentation, and predicts all-cause mortality. Furthermore, positive correlation of the SoM and the detrimental scores with the number of risk factors in an individual further suggest that each risk factor contributes to increased baseline dysregulation. However, even after adjusting for the effects f age, sex, BMI, disease and smoking statuses, the SoM score was significantly associated with all-cause mortality. We also note that the hazard ratio of the adjusted SoM score (HR=1.8) was substantially higher than a previously reported 54-gene signature, IMM-AGE, that captured immune aging (HR=1.05) [96]. This breadth of association despite the broad heterogeneity across thousands of samples from tens of independent studies strongly suggests the SoM signature has broader implications and has potential to enable assessing baseline immune health.

Our study has several limitations. First, we used predefined 42-gene signature [18], which in turn was identified using blood samples from patient with acute respiratory viral infections [20]. Reanalysis of these datasets using all genes may identify a larger set of genes involved in this conserved common immune response. On the other hand, the prior selection of the SoM gene signature may be useful as the number of hypotheses to be corrected is substantially lower. Second, majority of the data used in our analysis is from bulk transcriptome profiling of blood samples, which could be affected by changes in the proportions of immune cells or changes in transcriptome at a single-cell level. Although our analysis of >584,000 immune cells using scRNA-seq suggests changes in transcriptome, additional analyses are needed to investigate whether the changes in transcriptome at a single-cell resolution are markers of a change in proportion of specific immune cell subsets. Finally, similar to the IHM signature, the SoM quantifies system-wide dysregulation of immune response. However, we did not account for organ-specific immune responses that may vary depending on the organ involvement (e.g., respiratory tract in infection versus pancreas in diabetes). Further studies focusing on integrating peripheral esponses with organ-specific differences are required.

In summary, we performed one of largest multi-cohort transcriptome analysis with data spanning single-cell and bulk transcriptomics and bulk proteomics from 12026 samples across 68 studies and found the dysregulation between the detrimental and protective host responses defined by the SoM score to be associated with multiple risk factors for severe infection, bacterial infections and the differential treatment response to acute sepsis infection and chronic asthma. Our results point to the presence of a neutrophil subset in these conditions, which is characterized by increased expression of genes associated with detrimental host response. Our results provide testable hypotheses for future studies to uncover the mechanisms driving this increase in risk and design appropriate intervention strategies to alleviate this risk.

## Methods

### Collection, curation, and preprocessing of transcriptomic data for bacterial infection at presentation

We searched the National Center for Biotechnology Information (NCBI) Gene Expression Omnibus (GEO) and ArrayExpress for datasets comprised of samples from humans with bacterial infections, using search terms “sepsis”, “bacterial sepsis”, “bacterial infection”, terms for most specific organ system infections (e.g., urinary tract infection), and common infectious bacterial pathogens. We excluded datasets if they were missing healthy controls (required for COCONUT co-normalization), clinical severity information, confirmatory microbiological testing, or gene expression data for the genes in the SoM signature. We identified 17 datasets consisting of 828 blood samples from patients with bacterial infections (Table S1) [21–37].

We assigned one of seven clinical severity levels (healthy control, asymptomatic/convalescent, mild, moderate, serious, critical, or fatal) based on the patient’s symptoms, level of care, and dissemination of disease as previously described [18] (Figure 1A). Briefly, mild infections included patients with a minor focal infection that did not require inpatient care. Moderate infections included patients requiring admission to the hospital with a focal infection, or any systemic infection that did not require ICU-level care. Patients with serious infections were those admitted to the ICU without mechanical ventilation or inotropes, patients described as having “severe” infection without further specification, or inpatients requiring supplemental respiratory support outside of the ICU. Patients assigned to critical were those admitted to the ICU and required mechanical ventilation, inotrope support, or with a diagnosis of shock or central nervous system (CNS) involvement. Patients with detectable infection but resolution of symptoms were classified as convalescent and included with asymptomatic patients. Finally, we grouped patients with mild or moderate severity as “non-severe” and those with serious, critical, or fatal severity as “severe”.

We co-normalized gene expression data across datasets using COCONUT [97] (COmbat CONormalization Using conTrols) to eliminate batch effects. COCONUT co-normalization pools microarray data by correcting for batch effect using healthy controls and then applying this correction to non-control samples.

### Association of transcriptomic scores with response to treatment in the VANISH trial

It is known that steroids improve outcomes in severe COVID-19 infection, however studies have suggested that steroids may be harmful in subgroups of patients with sepsis. To evaluate whether detrimental and protective host response modules were associated with heterogeneity of treatment response to steroids, we performed a secondary analysis of the VANISH cohort. The VANISH trial was a multicenter, factorial-design, randomized, controlled clinical trial that evaluated vasopressin vs norepinephrine and hydrocortisone vs placebo in septic shock. A subset of these patients had genome-wide gene expression analysis performed that is publicly available (E-MTAB-7581). We hypothesized that patients with low SoM scores and high protective host response scores (i.e., lower risk patients) would have worse outcomes with steroid treatment. To evaluate this, we calculated the SoM module scores for each patient in VANISH. We then performed logistic regression evaluating the interaction term between scores and their component module scores (SoM, protective, detrimental) and steroid assignment with mortality, adjusting for age, sex, and the interaction between gene scores and APACHE score.

### Analysis of steroid effect on host-response at the bulk level

To evaluate whether differences in protective score could be detected at the bulk transcriptomic level, we used publicly available bulk transcriptome data from samples collected as part of the Low-dose Hydrocortisone in Acutely Burned Patients randomized clinical trial (GSE77791) [39, 40]. In short, this was randomized control study that randomized acute burn patients with vasodilatory shock to hydrocortisone 200mg/day versus placebo. Whole blood was collected from patients in the trial at baseline and 24-, 120-, and 168-hours post-randomization, and whole genome expression was measured. Gene signatures were calculated for all patients at all time points, and we evaluated the difference in gene scores between steroid and placebo-randomized patients at baseline and day seven using Wilcoxon rank sum test.

### Integration and visualization of single cell data

We collected and curated the single-cell blood transcriptome datasets from healthy controls and patients with COVID-19 or bacterial sepsis from four studies [42–45]. Datasets were processed with Seurat (v4.0.5) [98]. We removed cells with fewer than 50 detected genes, or fewer than 200 mRNA reads, or more than 20% mitochondrial reads. Datasets were integrated with reciprocal PCA algorithm in the Seurat integration workflow. Clusters were identified with Seurat SNN graph construction on PCA embeddings after integration, followed by a Louvain community detection algorithm. The cell types were annotated using canonical cell type markers. Cells were visualized in a low dimensional space using uniform manifold approximation and projection (UMAP).

### Collection and analyses of case control studies for analysis of risk factors

To identify studies profiling healthy subjects younger than 40 or older than 60, we searched NCBI GEO using search terms “vaccine response”, “healthy”, “adult”, and “old or elderly, young”. In total, we identified 8 datasets containing 375 blood samples [55–59] (Table S2). For sex, we used 16 datasets [60–73] we curated previously [2], containing 953 samples (Table S2). For obesity, we searched NCBI GEO using search terms “BMI”, “lifestyle”, “obese”, “lean”, “healthy”, and “adult”. We excluded datasets where the definition of lean and obese were different from BMI*≤*25 and BMI*≥*30 respectively. In total, we identified 4 datasets containing 180 samples [74–77] (Table S2). For diabetes, we searched NCBI GEO using search terms “diabetes”, “adult”, “blood”. In total, we identified 10 datasets containing 313 samples [78–85] (Table S2). For all risk factors, we excluded datasets where there were fewer than 5 cases or controls. We used MetaIntegrator [99, 100], our multi-cohort meta-analysis framework, to compute summary effect sizes and p-values for each risk factor.

### Processing of data from Framingham Heart Study

We used data from the Offspring and Generation III cohorts in the Framingham Heart Study [63], since only those two cohorts contained gene expression measurement. We only used the subjects in the consent group 1 denoted c1.HMB-IRB-MDS.

For gene expression measurement, we used data as deposited in v18 in phe000002. The gene expression for the Offspring and Generation III cohorts, measured from whole blood samples taken during exam 8 and 2 respectively, were taken from FinalFile_Gene_OFF_2446_Adjusted_c1.txt and Fina_Exon_GENIII_3180_c1.txt respectively. We used the Affymetrix platform table GPL5188.txt to process the expression data. We conormalized the expression from the two cohorts using COCONUT [97], using the healthy subjects from both the cohorts as the reference. We computed the scores for SoM, detrimental, protective, and modules 1-4 on the conormalized data.

We used the phenotypic and clinical measurements from data deposited in v31, corresponding to exams 8 for Offspring and 2 for Generation III cohorts. We used age at the time of gene expression measurement and sex from the table pht003099. Using data from pht000309, we identified patients with cardiovascular events as those having an EVENT corresponding to {1, 2, 3, 4, 5, 6, 7, 10, 11, 12, 13, 16, 17, 18, 21, 22, 23, 24, 25, 26, 30, 39, 40, 41}. We denoted patients with cancer as those having an entry in pht000039. We identified patients with current diabetes from pht000041 for the Offspring cohort and pht007775 for the Generation III cohort. Using data from pht000691, we denoted patients with dementia as those with mri352 values different than 0. Using data from pht000747 for the Offspring cohort and pht003094 for the Generation III cohort, which contained questionnaires, doctor impressions, and clinic exam taken at the time of gene expression measurement, we identified height, weight, and whether a subject had any fever or illness in the past two weeks. Using these tables, we also identified if a subject had any potential illnesses or diseases based on doctor impression or questionnaires if the values of any of {H339, H340, H341, H342, H344, H345, H347, H348, H349, H354, H357, H358, H359, H360, H363, H364, H367, H368, H369, H370, H374, H375, H378, H379} in the Offspring cohort or {g3b0404, g3b0405, g3b0406, g3b0408, g3b0407, g3b0410, g3b0411, g3b0414, g3b0415, g3b0416, g3b0418, g3b0420, g3b0422, g3b0424, g3b0427, g3b0429, g3b0430, g3b0431, g3b0432, g3b0434, g3b0435, g3b0436, g3b0439, g3b0441, g3b0442, g3b0444, g3b0445, g3b0452, g3b0453, g3b0454, g3b0448, g3b0449, g3b0426, g3b0446} in the Generation III cohort was 1. We identified current or former smokers as those who had either of H060, H066 in the Offspring cohort or g3b0091, g3b0097 in the Generation III cohort to be different than 0. Using the same pair of measurements, we identified those that quit smoking as having the first measurement to be 1 and the second measurement to be a positive integer. We identified healthy subjects as those that did not have any cardiovascular event, cancer, diabetes, dementia, fever or illness in the past two weeks, potential illnesses or diseases based on doctor impression or questionnaires. Lastly, we identified mortality events using pht003317.

### Calculation of the IHM score

The immune health metric (IHM) score [88] is computed as the difference in the geometric means of the expression between *{SLC16A10, NDRG2, AK5, CD7, RNFT2, PHC1, MAN1C1, FAM102A, RCAN3, EIF3H, RPS17, RPS5, PLXDC1, TESPA1, SGK223, EIF3L, ANAPC16, CD27, NPAT, ID3, RACK1, APEX1, GCNT4, FCMR, TCF7, KLHL3, AXIN2, LY9, RPS25, LDHB, PKIA, RPL3, N6AMT1, GAL3ST4, SSBP2, CD1C, LEF1, RPL7, PIK3IP1, GPRASP1, ABI2, APBB1, SPTBN1, GPA33, CCR9, BCKDHB, SCAI, RPL4, NOG, TCEA3, ETS1, LDLRAP1, GPR183, ZNF548, ZNF91, NPM1, MSANTD2, KAT6B, SLC7A6, DCHS1, OXNAD1, RPS2, RPL7A, GRAP, RPL23A, RPL10A, RPS3, FAM175A, RPL29, EEF2, EDAR, ABLIM1, MBLAC2, CCR7, ZNF573, CAMK4, LRRN3, MAGI3, RPLP2, ZIK1, NT5E, FUT8, ZNF101, RPL34, RPS20, FOXP1, ZNF550, TSPYL2, ATP6V0E2-AS1, GRPEL2, MGC57346, SEPT1, PRKACB, AGMAT, RPL11, MYC, ZZZ3, RPL5 }* and *{CTRL, NTNG2, AP5B1, PDCD1LG2, DOCK4, S100A9, BACH1, FAM8A1, SECTM1, S100A8, TYMP, HK3, IL1RN, MYD88, REC8, ALPK1, SAT1, PRKCD, SLC26A8, PARP9, LMNB1, RELT, TAP1, JAK2, BRI3, GBP2, PLEKHO2, ETV7, ODF3B, SIGLEC5, CEACAM1, CARD16, ZBP1, DDX60L, APOL2, CD63, TNFAIP6, KCNJ2, ANKRD22, SCARF1, SEMA4A, DNAJC5, SQRDL, HELZ2, GADD45B, FAS, HSPA6, PIK3AP1, CLEC7A, SERPING1, OR52K2, ITPRIP}*. The IHM score is designed to be higher in healthy individuals whereas the SoM is designed to be lower in healthy individuals.

### Description of the Arivale cohort

The Arivale cohort was derived from 6,223 individuals who participated in a wellness program offered by a currently closed commercial company (Arivale Inc., Washington, USA) between 2015–2019. An individual was eligible for enrollment if the individual was over 18 years old, not pregnant, and a resident of any U.S. state except New York; participants were primarily recruited from Washington, California, and Oregon. The participants were not screened for any particular disease. Plasma concentrations of proteins in the Arivale cohort were measured using the ProSeek Cardiovascular II, Cardiovascular III, and Inflammation panels (Olink Biosciences, Uppsala, Sweden) at Olink facilities in Boston, MA. The ProSeek method is based on the highly sensitive and specific proximity extension assay, which involves the binding of distinct polyclonal oligonucleotide-labelled antibodies to the target protein followed by quantification with real-time quantitative polymerase chain reaction (rt-PCR) [101]. Samples were processed in several batches; potential batch effects were adjusted using pooled control samples included with each batch. In this study we limited the original cohort to the participants whose datasets contained baseline proteomics. This study was conducted with de-identified data of the participants who had consented to the use of their anonymized data in research. All procedures were approved by the Western Institutional Review Board (WIRB) with Institutional Review Board (IRB) (Study Number: 20170658 at Institute for Systems Biology and 1178906 at Arivale).

## Acknowledgements

P.K. is funded in part by the Bill and Melinda Gates Foundation (OPP1113682); the National Institute of Allergy and Infectious Diseases (NIAID) grants 1U19AI109662, U19AI057229, and 5R01AI125197; Department of Defense contracts W81XWH-18-1-0253 and W81XWH1910235; and the Ralph & Marian Falk Medical Research Trust. The funders had no role in study design, data collection and analysis, decision to publish, or preparation of the manuscript.

## Author contributions

A.G., A.R.M., H.Z., J.T., and P.K. conceived the study. P.K. supervised the study. A.T.M. and J.R.H. processed data from the Arivale cohort. A.G., A.R.M., H.Z., J.T., and M.F. collected, annotated, processed, and analyzed data. A.G., A.R.M., H.Z., and P.K. interpreted analysis results and wrote the manuscript.

## Supplementary tables and figures

**Table S1.** Summary of datasets used for evaluating the association between the SoM score and severity of bacterial infections.

| Dataset | Publication | Tissue | Cases | Controls | Females |
| --- | --- | --- | --- | --- | --- |
| EMTAB3423 | Waddington *et al.*, CID, 2014 | Whole blood | 119 | 40 | 38 |
| GSE103119 | Wallihan *et al.*, Front Cell Infect Microbiol, 2018 | Whole blood | 35 | 20 | 25 |
| GSE113867 | Blohmke *et al.*, EMBO Mol Med, 2019 | Whole blood | 81 | 135 | 63 |
| GSE128557 | Ambite *et al.*, PLoS Pathog, 2019 | Whole blood | 4 | 5 | 4 |
| GSE145974 | Petzke *et al.*, mBio, 2020 | Peripheral blood mononuclear cells | 65 | 21 | 43 |
| GSE16129 | Ardura *et al.*, PLoS One, 2009 | Peripheral blood mononuclear cells | 55 | 19 | 29 |
| GSE16463 | Tantibhedhyangkul *et al.*, PLoS Negl Trop Dis, 2011 | Peripheral blood mononuclear cells | 11 | 2 | 9 |
| GSE25504 | Smith *et al.*, Nat Commun, 2014 | Whole blood | 37 | 44 | 31 |
| GSE30119 | Banchereau *et al.*, PLoS One, 2012 | Whole blood | 99 | 44 | 66 |
| GSE46955 | Shalova *et al.*, Immunity, 2015 | Monocytes | 32 | 12 | 10 |
| GSE47172 | Kulohoma *et al.*, BMJ Paediatr Open, 2017 | Whole blood | 6 | 3 | 4 |
| GSE60244 | Suarez *et al.*, J Infect Dis, 2015 | Whole blood | 22 | 40 | 33 |
| GSE6269 | Ramilo *et al.*, Blood, 2007 | Peripheral blood mononuclear cells | 73 | 6 | 33 |
| GSE64456 | Mahajan *et al.*, JAMA, 2016 | Whole blood | 89 | 19 | 49 |
| GSE68004 | Jaggi *et al.*, PLoS One, 2018 | Whole blood | 17 | 37 | 29 |
| GSE72946 | Lindow *et al.*, PLoS Pathog, 2016 | Whole blood | 29 | 4 | 3 |
| GSE9960 | Tang *et al.*, Crit Care Med, 2009 | Peripheral blood mononuclear cells | 54 | 16 | 27 |

**Table S2.**
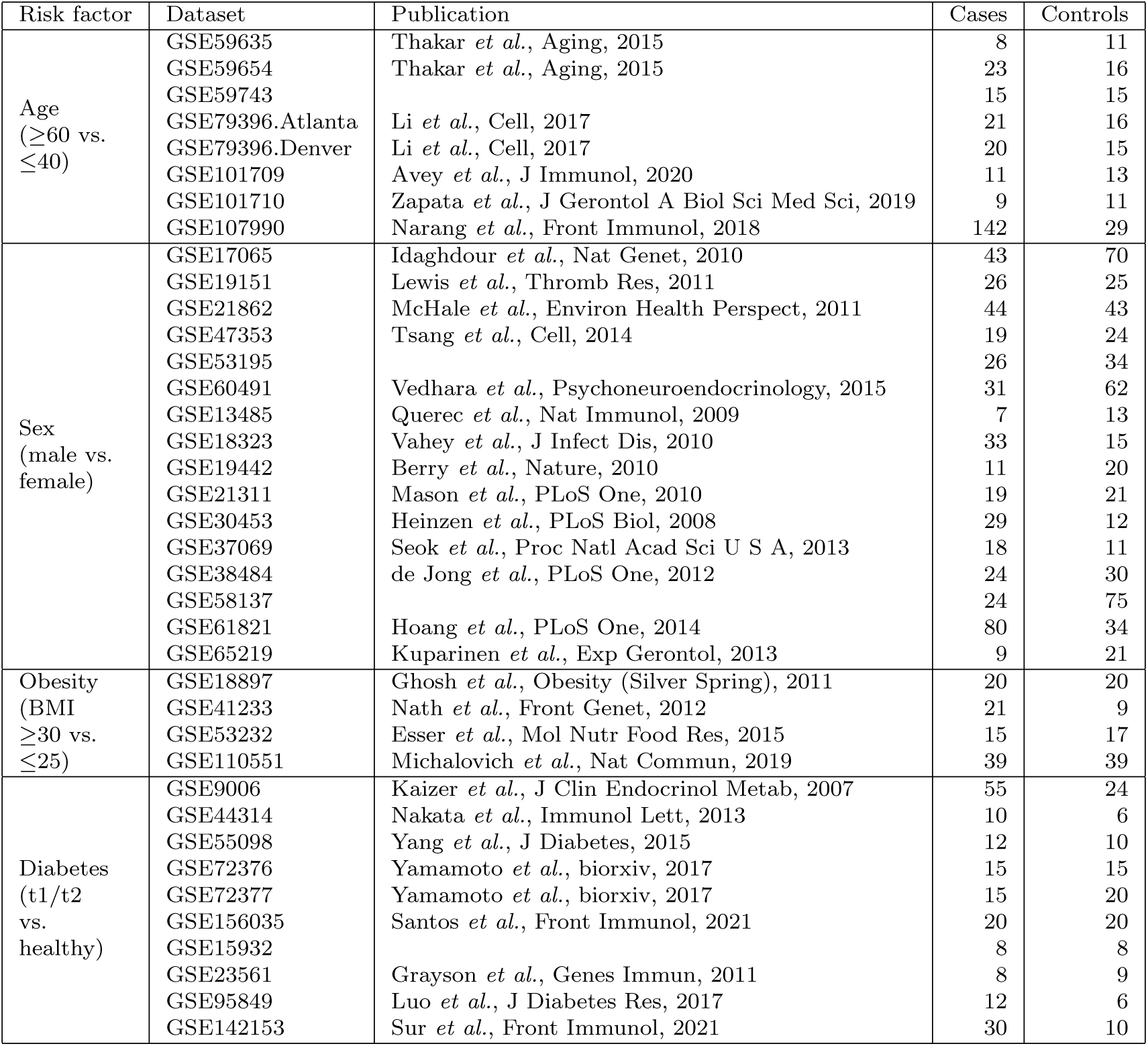
Summary of datasets used for case-control meta-analyses for evaluating the association between the SoM score and risk factors for severe infections.

**Fig. S1.**
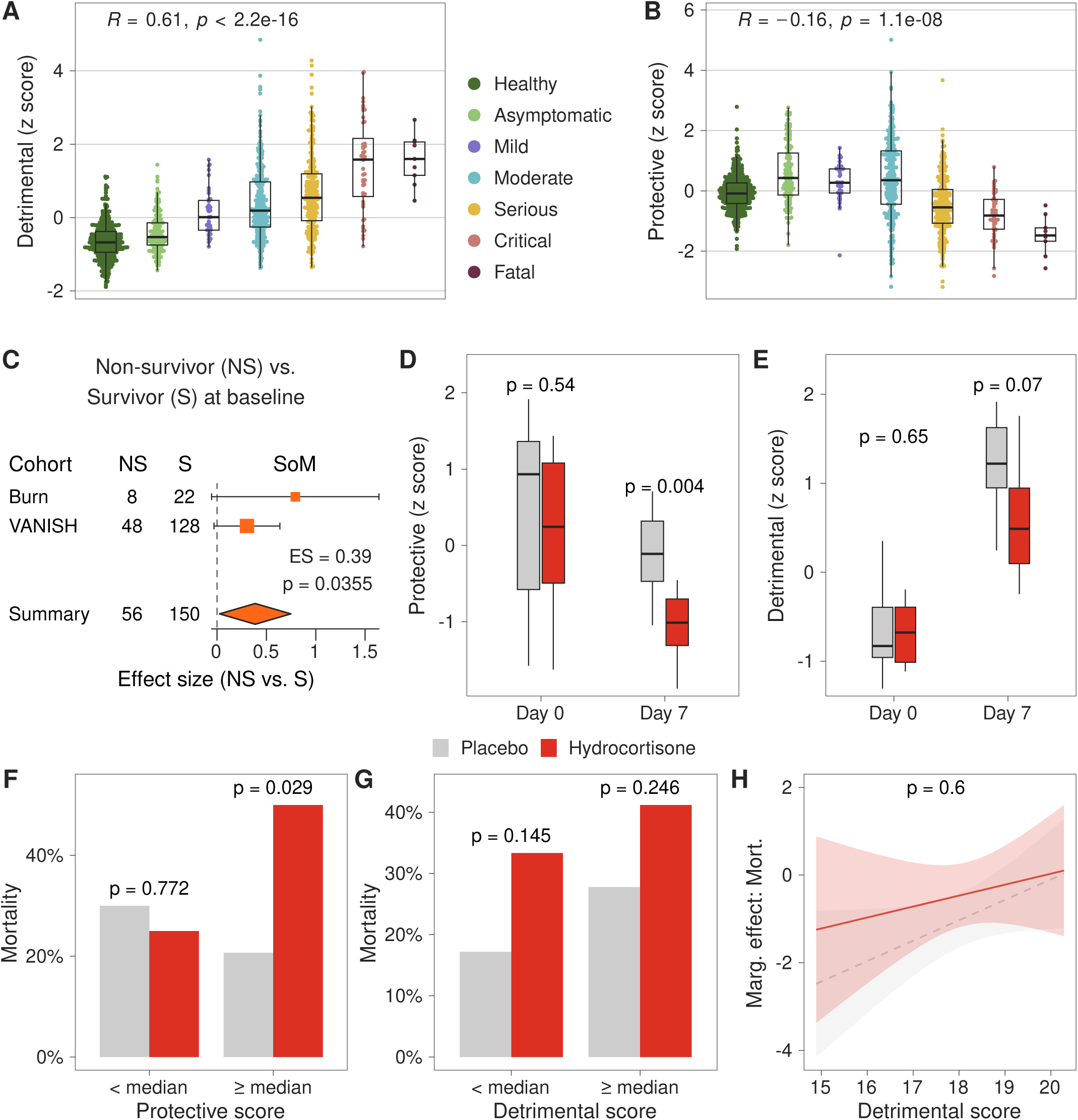
**(A-B)** Variation of the z-scaled (**A**) detrimental, and (**B**) protective scores with severity of bacterial infection. R denotes Pearson correlation, and p its corresponding p-value. (**C**) Forest plot with effect size for the SoM score between non-survivors (NS) and survivors (S) in the Burn and VANISH cohorts. Summary effect size and p-value were computed using a random effects model. (**D-E**) Box plots of z-scaled (**D**) protective and (**E**) detrimental score comparing placebo and hydrocortisone arms on days 0 and 7 in the Burn cohort. P-values were calculated using the Wilcox test. (**F**) Bar plot of mortality comparing the randomized placebo and hydrocortisone arms for subjects with baseline protective scores lower than and higher than median in the Vanish cohort. P-values were calculated using the Fisher test. (**G**) Bar plot of mortality comparing the placebo and hydrocortisone arms for subjects with baseline detrimental scores lower than and higher than median in the Vanish cohort. P-values were calculated using the Fisher test. (**H**) Interaction plot showing the marginal dependence of mortality on the detrimental score for the placebo and hydrocortisone arms in the Vanish cohort. Logistic model p-value for the interaction term between detrimental score and steroid arm is shown.

**Fig. S2.**
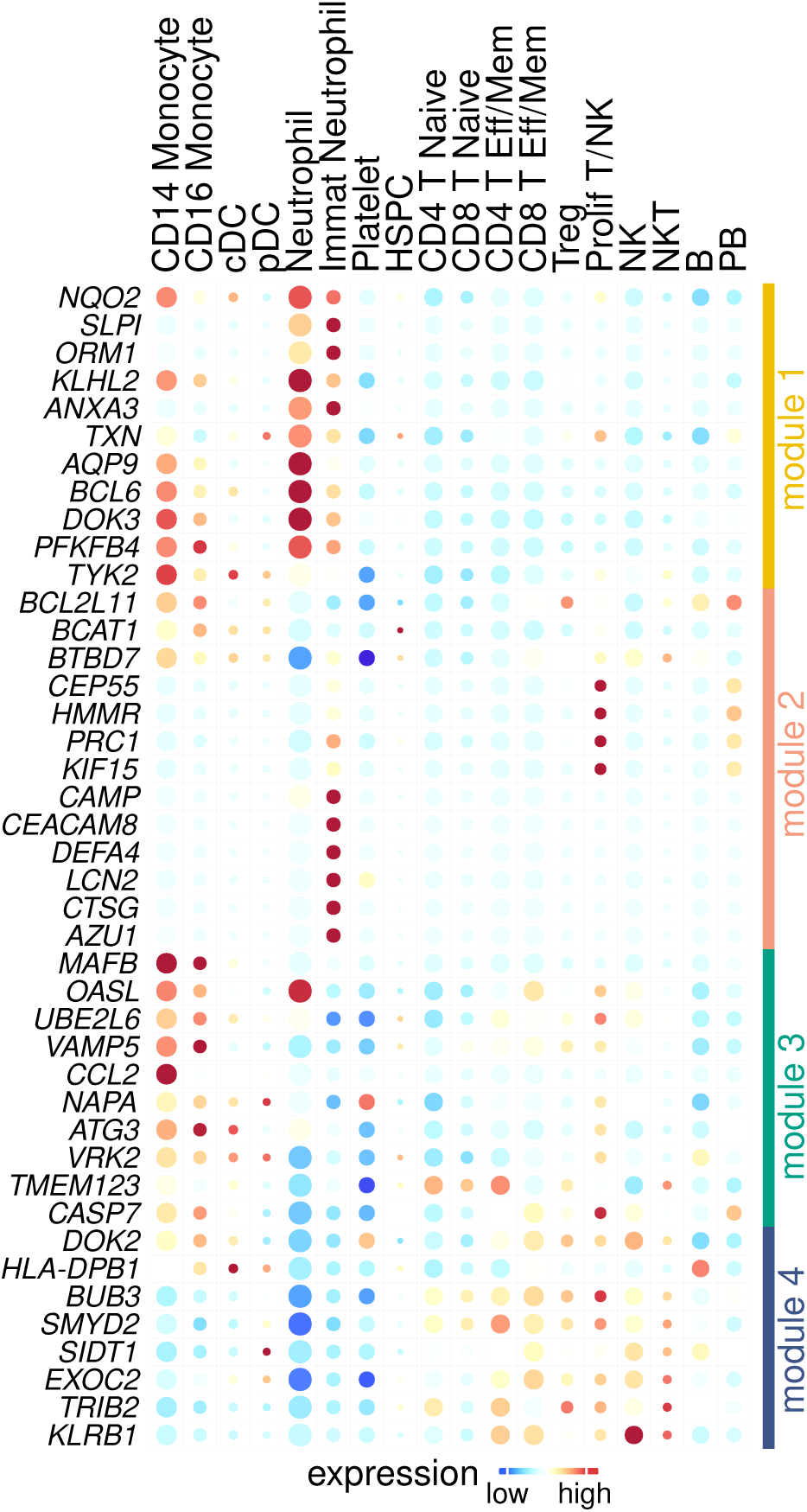
Heatmap of average expression the 42 SoM genes in 20 cell types. Size of the circle is proportional to the cell type proportion.

**Fig. S3.**
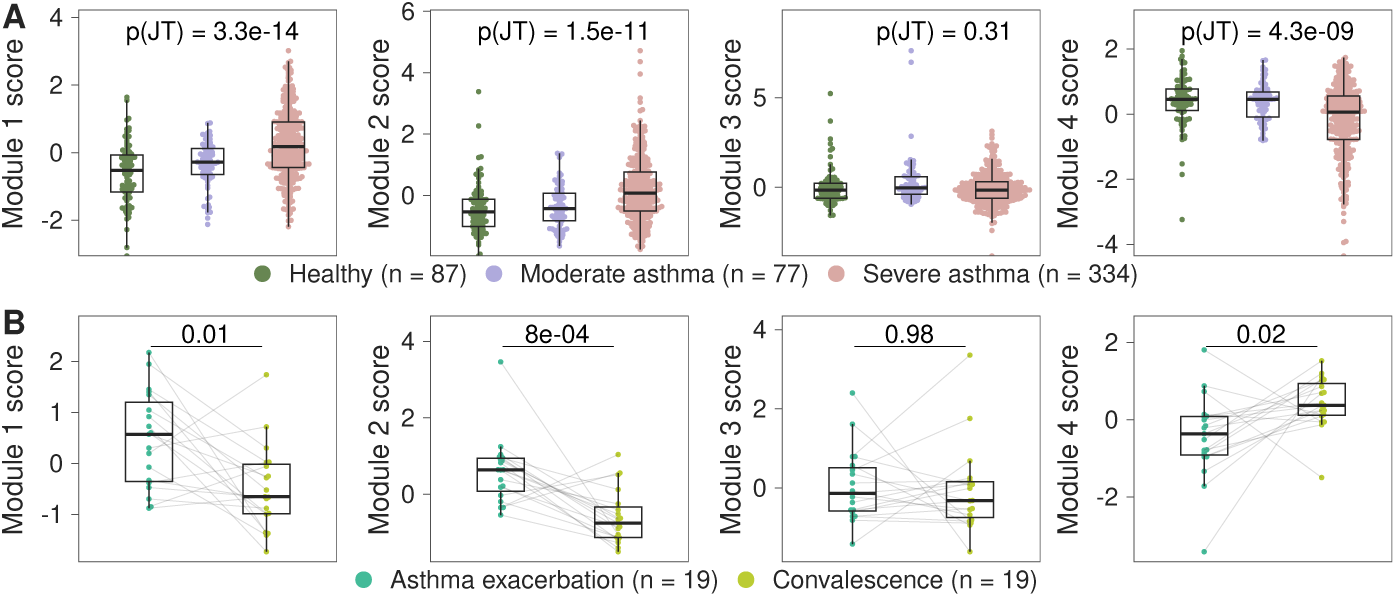
(**A**) Variation of the z-scaled module 1, 2, 3, and 4 scores with asthma severity. P-values were calculated using the Jonckheere–Terpstra trend test. (**B**) Comparison of the z-scaled module 1, 2, 3, and 4 scores at asthma exacerbation and after convalescence. P-values were calculated using the paired Wilcox test.

**Fig. S4.**
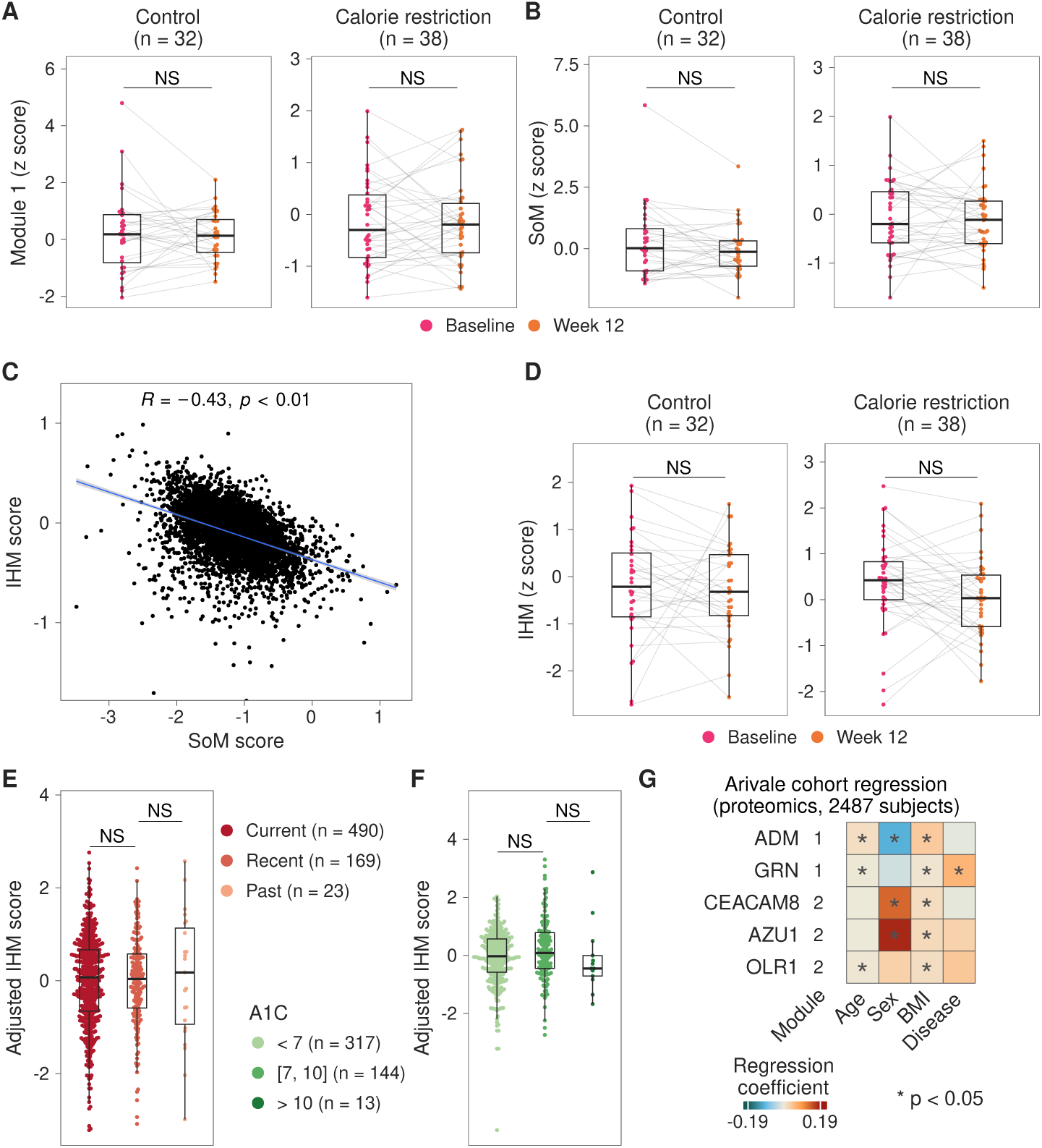
(**A, B, D**) Comparison of the z-scaled (**A**) SoM, (**B**) module 1, and (**D**) immune health metric (IHM) scores at baseline and after 12 weeks for the control and calorie restriction groups in a randomized diet restriction study. P-values were computed using the paired Wilcox test. (**C**) Scatter plot between the IHM and SoM scores for each subject in the Framingham cohort. R denotes Pearson correlation, and p its corresponding p-value. (**E-F**) Variation in the IHM score adjusted for age, sex, BMI and (**E**) disease or (**F**) smoking status among (**E**) current, recent (quit within the last 5 years), and past (quit more than 5 years ago) smokers and (**F**) subjects with controlled (A1C<7), moderately uncontrolled (7*≤*A1C*≤*10), and highly uncontrolled (A1C>10) diabetes. P-values were computed using the Wilcox test. (**G**) Summary of linear regression models for risk factors in the Arivale proteomic cohort. Heatmap color depicts the coefficient of regression and the circle size the corresponding p-value for a given protein abundance/module score and risk factor. * denotes p<0.05.

**Fig. S5.**
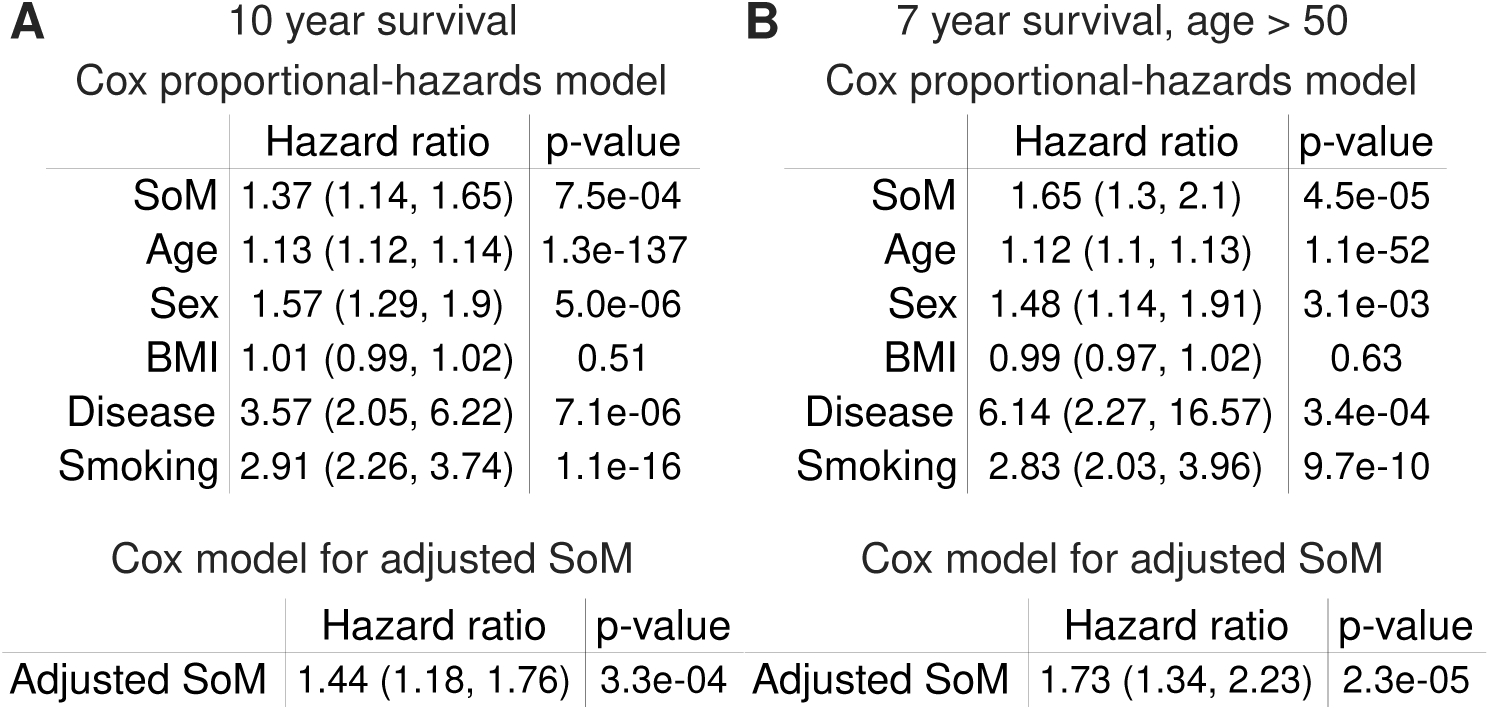
Summary of Cox proportional-hazards models in the Framingham cohort evaluating the association with (**A**) 10-year mortality and (**B**) 7-year mortality among subjects older than 50 years for the SoM score, age, sex, BMI, disease and smoking statuses, and the SoM score adjusted for age, sex, BMI, disease and smoking statuses. Hazard ratio, its 95% confidence estimates, and the corresponding p-values are shown.

